# TissueGrinder, a novel technology for rapid generation of patient-derived single cell suspensions from solid tumors by mechanical tissue dissociation

**DOI:** 10.1101/2021.06.08.447529

**Authors:** Stefan Scheuermann, Jonas M. Lehmann, Ramkumar Ramani Mohan, Christoph Reißfelder, Felix Rückert, Jens Langejürgen, Prama Pallavi

## Abstract

**Introduction:** Recent advances hold promise of making personalized medicine a step closer to implementation in clinical settings. However, traditional sample preparation methods are not robust and reproducible. In this study, the TissueGrinder, a novel mechanical semi-automated benchtop device, which can isolate cells from tissue in a very fast and enzyme-free way is tested for cell isolation from surgically resected tumor tissues.

**Methods:** 33 surgically resected tumor tissues from various but mainly pancreatic, liver or colorectal origins were processed by both novel TissueGrinder and explant method. An optimized processing program for tumors from pancreatic, liver or colorectal cancer was developed. The viability and morphological characteristics of the isolated cells were evaluated microscopically. Expression of pancreatic cancer markers was evaluated in cells isolated from pancreatic tumors. Finally, the effect of mechanical stress on the cells was evaluated by assessing apoptosis markers via western blotting.

**Results:** TissueGinder was more efficient in isolating cells from tumor tissue with a success rate of 75% when compared to explant method 45% in terms of cell outgrowth six weeks after processing. Cells isolated with TissueGinder had a higher abundance and were more heterogeneous in composition as compared to explant method. Mechanical processing of the cells with TissueGrinder does not lead to apoptosis but causes slight stress to the cells.

**Discussion:** Our results show that TissueGrinder can process solid tumor tissues more rapidly and efficiently and with higher success rate compared to the conventionally used explant method. The results of the study suggest that the TissueGrinder might be a suitable method for obtaining cells, which is important for its application in individualized therapy. Due to the great variance in different tumor entities and the associated individual tissue characteristics, a further development of the dissociation protocol for other types of tumors and normal will be targeted.

## 1 Introduction

Cancer is currently one of the most frequent causes of death in the western world and its prevalence is constantly increasing in absolute numbers [1, 2]. Although we have seen significant improvement in the line of treatment during the last decades, yet they proved to be adequate only in a number of patients [3, 4]. This is not only burdensome for the patient but also stresses healthcare systems. As every tumor shows a distinct biological heterogeneity, personalized cancer therapy seems to be the only reasonable treatment modality [5, 6]. Patient tumor derived cells can serve as avatars on which new treatments, such as chemotherapeutics, or hypotheses for the improvement of treatments can be tested for effectiveness prior to administering drugs to the patient, thus providing better choices in the hands of physicians and minimizing adverse effects for the patients [7–9].

The combination of personalized medicine with the increasing availability of novel diagnostic technologies has enormously increased the demand for sufficient quantities of high-quality biological samples [3, 10–18]. Many personalized medicine approaches involving single cell sequencing or drug screening rely on cell isolation from biopsy or surgically resected tumor material [8, 19–21]. In addition, clinical tissue samples for the obtaining primary cells are often only available in small quantities. While analytical methods (e.g. Next Generation Sequencing, Flow Cytometry, single cell DNA/RNA sequencing and high throughput screening) have become considerably faster and more sensitive in recent years, this does not apply to the sample preparation. These tissue-processing workflows done prior to the actual analysis are still time-consuming processes as these are often performed manually, and are not completely reproducible [15].

To obtain primary cell cultures, explant and tissue dissociation procedures (mechanical and enzymatic) are generally used [17, 22, 23]. The explant method dominated the field of tissue culture for more than 50 years to obtain primary cells [24, 25]. This is one of the simplest procedures, briefly, tissue explant is finely chopped into fragments (1 to 2 mm3) and placed in a culture flask [26]. Cells migrate out of the explants and grow on the culture surface. However, success has been limited by long processing times, low yield and high manual workload leading to a significant increase in the overall time from tissue resection to obtaining a cell line [27]. Apart from the explant method, enzymatic, chemical and mechanical dissociation are three methods widely used for tissue dissociation [28]. The dissociation with chemical or enzymatic reagents is not always desired because they also affect the proteins that might be later needed for labeling or staining, or are required for molecular biological analysis [17]. In the mechanical approach, no additional enzymes or other agents are used to isolate cells from tissue, which could potentially alter the cells. However, with mechanical dissociation, the cells undergo some mechanical stress.

TissueGrinder technology facilitates isolation of individual cells from biological tissue via mechanical dissociation where mechanical stress can be reduced to a minimum in order to obtain cells that are as identical as possible to the body, so that diagnostics done on these cells are more effective. This technology applies a combination of shearing and cutting forces to dissociate cells from precut tissue (Figure 1 B) [29]. The duration and speed of shearing and cutting forces can be optimized to develop tissue specific dissociation protocols. Manually diced tissue sample is transferred into grinding tubes, which contains an integrated grinding gear (Figure 1A). After dissociation, centrifugation of the grinding tubes, allow easy collection of cells which can be either cultured directly or used for further analytical tests. Thus, TissueGrinder combines tissue specific dissociation protocols with gentle mechanical dissociation to provide high quality single cell suspensions without use of enzymes.

**Figure 1:**
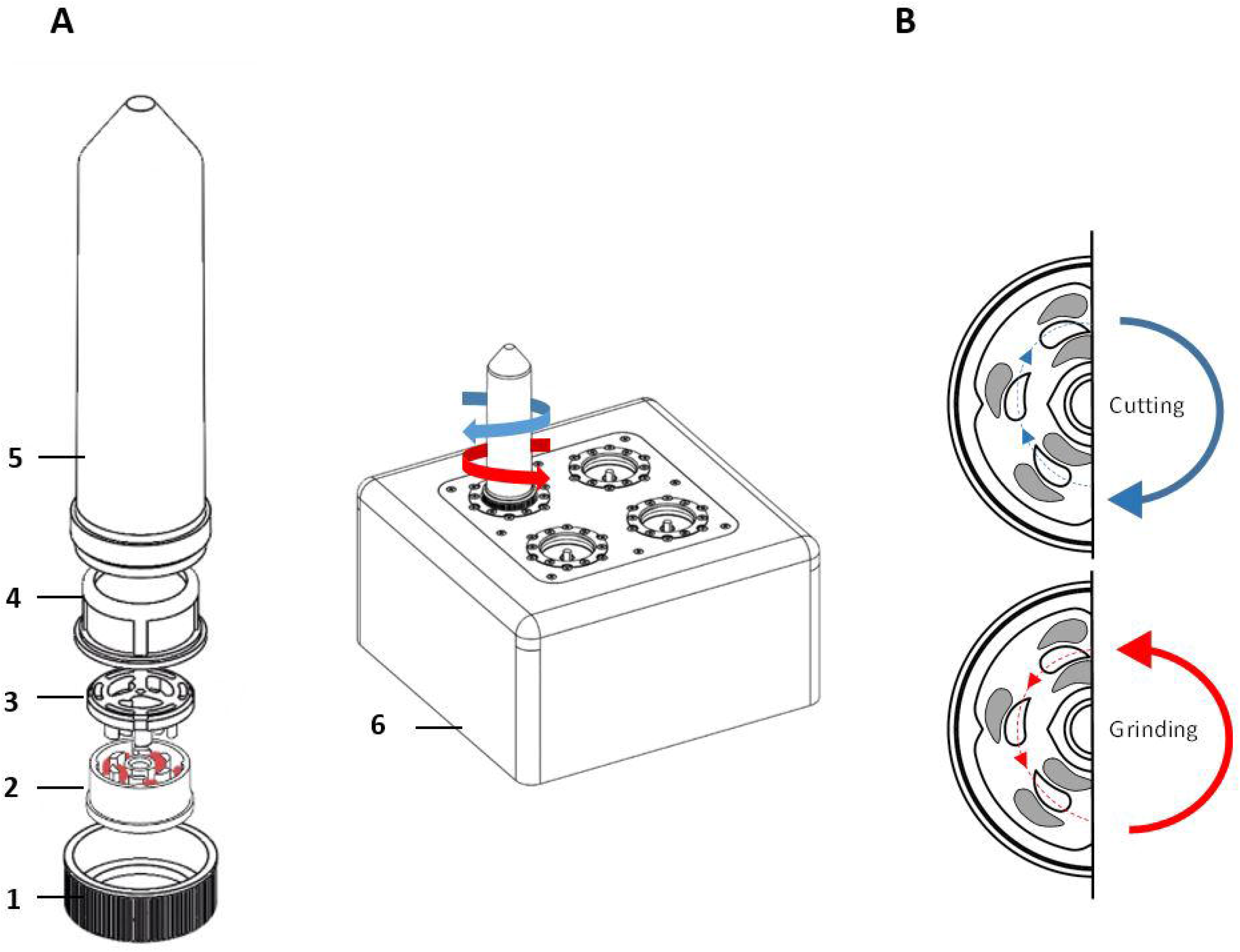
Assembly of single use Tissue Grinder sterile grinding set. A) The grinding set consists of a fitting lid to the 50 ml centrifuge tube (Falcon Tube) with a centered hole (1), the rotor (2) and the stator (3) both forming the grinding gear, a standard cell strainer with a pore size of 100 μm (4), and a 50 mL centrifuge tube (Falcon Tube)(5). The assembled grinding set is locked with the rotor’s specific notch to its counterpart on the TissueGrinder (6). The diced tumor tissues, each with edge lengths of 1-2 mm and a total weight of 100-200 mg, were placed in the inner part of the rotor and the unit was assembled as shown under a cell culture bench. B) Diced tissue fragments are processed into single cells by alternating processes of grinding and cutting which is achieved in the TissueGrinder by clockwise or anticlockwise turning of the rotor.

Due to the high demand for primary cells and the limited availability of tissue samples, the isolation of primary cells should be as efficient as possible [30]. In this study, we develop and describe a workflow for generation of cancer patient derived primary cell cultures using TissueGrinder. The developed workflow was compared with the standard explant method. These methods were analyzed according to the following parameters: sample preparation time, cell isolation efficiency, time until cells start to grow and heterogeneity of isolated cells. In addition to this we also evaluate if the stress caused by mechanical dissociation led to apoptosis in the isolated cells.

## 2 Methods

Surgically resected tumor tissues were collected and processed within one hour by two different techniques for isolation of the cells – explant method and by using Tissue Grinder. This study was approved by the local ethics committee of the University Medical center Mannheim (2012-293N-MA), and all of the donors gave their written, informed consent.

### 2.1 Tissue preparation

The surgically resected tumor tissue was transported on ice from operation room to laboratory and processed immediately. Tumor tissue was washed with PBS supplemented with 1% Penicillin/Streptomycin solution (Penicillin (10,000 units/mL) and Streptomycin (10,000 μg/mL)-solution, Gibco, Life Technologies Limited, USA). Blood-soaked tissue samples were washed with an ammonium chloride solution (Stemcell Technologies, Canada) to lyse present erythrocytes. Afterwards, the tissue was cut into small cubes with an edge length of about 1 mm. Subsequently, the tissue pieces were weighted and an equal amount of the tissue was processed for the cell isolation by explant method and Tissue Grinder.

### 2.2 Cell isolation via explant method

For explant cultures, the weighed, diced tumor tissue was placed in six well culture plate with 3 mL Dresden-Medium for at least six weeks to facilitate cell outgrowth. The culture was maintained at 37°C with 5% CO_2_ in a humidified atmosphere. Medium was changed every third day and culture plates were observed for cells growth. If no cell growth was observed after six weeks of culture, the culture plates were discarded.

### 2.3 Cell isolation using Tissue Grinder

For processing of the tissue with the TissueGrinder, the disposable sterile tube sets for the TissueGrinder were opened and assembled by latching the stator into the cell sieve under the cell culture bench. Afterwards, the cell sieve was transferred to the 50 mL centrifuge tubes. Stator and cell sieve were washed with 1 mL of Dresden-Medium. 100-200 mg of the prepared tissue pieces were transferred in the inner circle of the rotor (using a 1 mL pipette). 200μl of the medium was added to the tube. Finally, the rotor was placed on the stator in the cell sieve and covered with the lid with a centered hole. The assembly of the grinding unit is shown in Figure 1 A.

Afterwards, the closed tube is fixed on a port on the Tissue Grinder. Initially eight samples (MCC-190, MaPac-172, SFT, Carcinoma of unknown primary, MaPaC-174, MaPac-175, HCC-183 and CRC-12) were processed with three different programs – soft, medium and hard described in Table 1 to obtained right program for isolation of the cells.

**Table 1:**
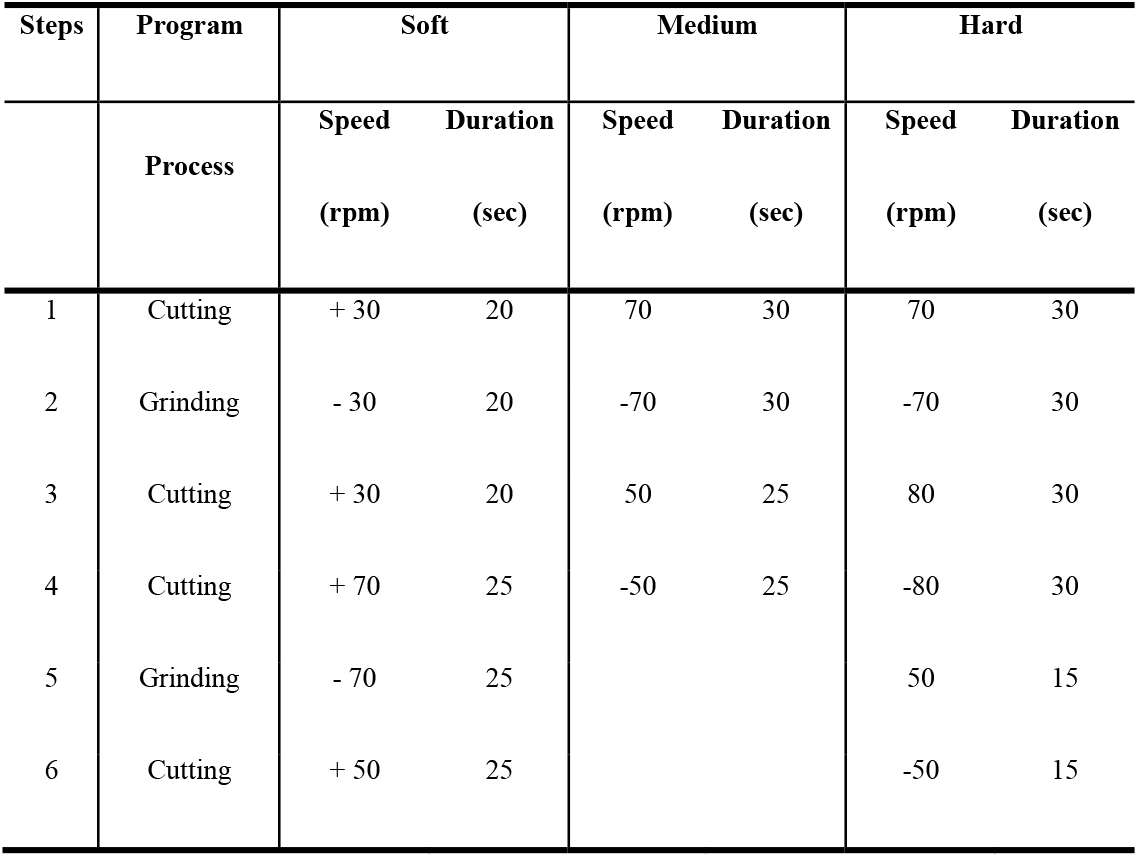
Table describing steps of the TissueGrinder programs used for processing of tumor tissue. It shows the two different process – cutting and grinding which are repeated alternatively along with the speed and duration for each of the tissue processing programs. A positive speed means that the TissueGrinder turns clockwise which leads to a cutting of the tissue. Whereas a negative speed symbolizes that the TissueGrinder turns anticlockwise which causes a grinding of the tissue. Before each step, the TissueGrinder takes 5 sec to adjust to the upcoming change. As a result, the duration of the whole program sums up to 3 min.

| Steps | Program | Soft |  | Medium |  | Hard |  |
| --- | --- | --- | --- | --- | --- | --- | --- |
|  | Process | Speed<br>(rpm) | Duration<br>(sec) | Speed<br>(rpm) | Duration<br>(sec) | Speed<br>(rpm) | Duration<br>(sec) |
| 1 | Cutting | + 30 | 20 | 70 | 30 | 70 | 30 |
| 2 | Grinding | - 30 | 20 | -70 | 30 | -70 | 30 |
| 3 | Cutting | + 30 | 20 | 50 | 25 | 80 | 30 |
| 4 | Cutting | + 70 | 25 | -50 | 25 | -80 | 30 |
| 5 | Grinding | - 70 | 25 |  |  | 50 | 15 |
| 6 | Cutting | + 50 | 25 |  |  | -50 | 15 |

The program drives the cutting and grinding processes in alternating fashion and varying intensities. Following the run, the tube was centrifuged for 5 minutes with 350 g. After centrifugation the tube was opened again in the laminar flow cabinet. Tissue remains were flushed with 1 mL Dresden-Medium into the cell sieve to ensure that all cells were collected in the tube. The tissue fragments, which did not pass through the sieve were discarded. The cell sieve including the stator was removed from the tube. The tube was closed by a 50 falcon lid without a hole and centrifuged for 5 minutes at 350 g. The cell pellet was resuspended with 3 mL Dresden-Medium and cultured in a separate well of a six well plate for each run. The complete workflow is depicted in Figure 2.

**Figure 2:**
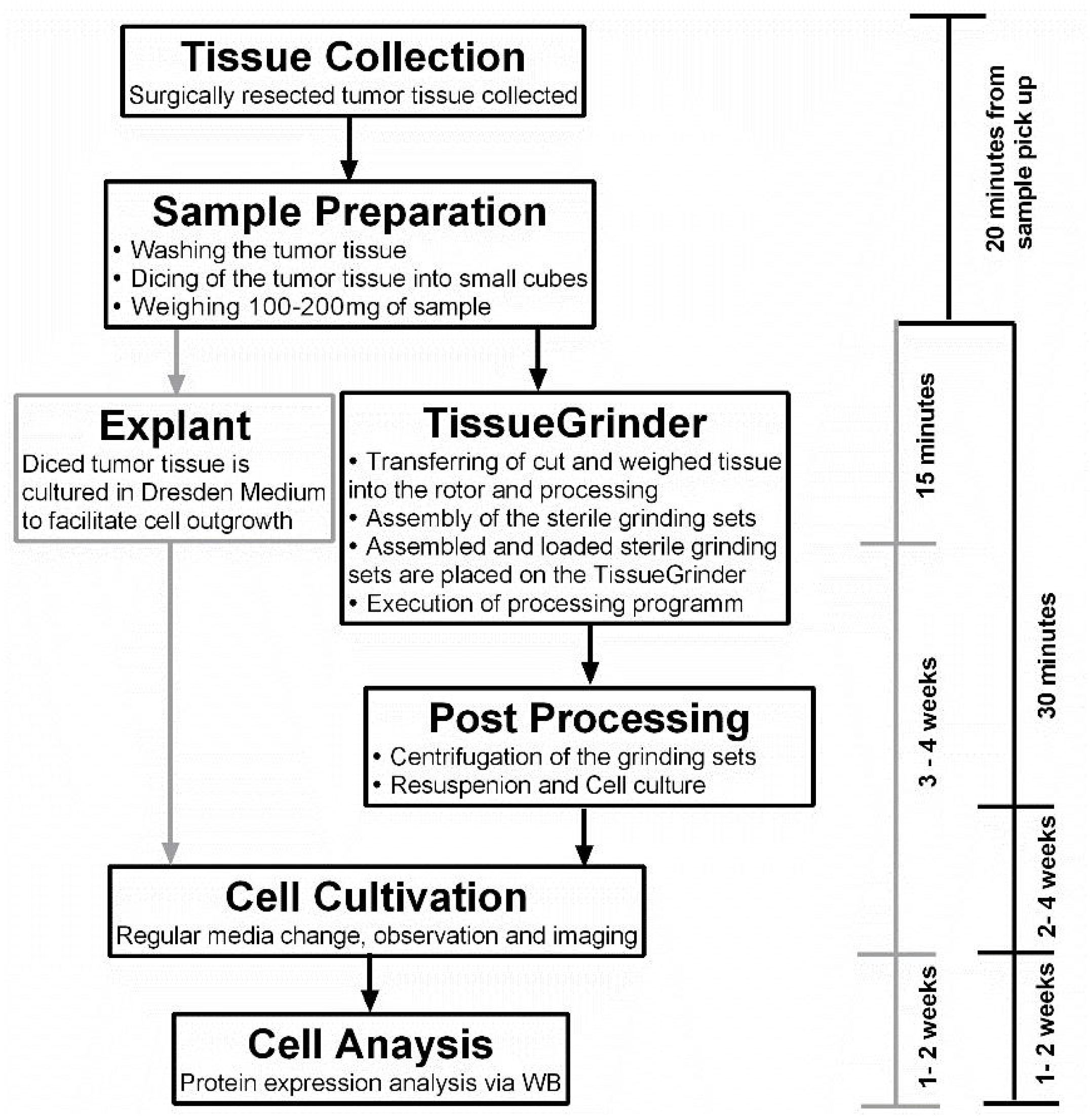
The complete workflow of tumor tissue processing. The tissue was collected as soon as possible from the operation room and the sample preparation started within 1 h of the resection of the tumor. The sample preparation time was 15-30 minutes for the TissueGrinder as well as for the explant method which is used as control. This included washing, cutting and weighing of tumor tissue. The TissueGrinder was assembled as shown in Figure 1 including 100-200 mg of tumor tissue that was placed in the inner part of the rotor. After closing the set, it was placed on the TissueGrinder and the program shown in Table. 1 was executed. Afterwards the tube was centrifuged, the supernatant discarded, the cell pellet resuspended and cultured in a separate well on the same six well plate as used for the explant method. Medium was changed every three days and cell growth was observed and documented using Zeiss Axiovert100 Microscope.

### 2.4 Cell Culture

Obtained cells were cultured in a 2D monolayer on six well plates until confluent, and sub-cultured into T25- and T75-Flasks. All cells received “Dresden-Medium” which is a mixture of two thirds of the common DMEM (Sigma-Aldrich Chemie GmbH, Steinheim, Germany) supplemented with 10% FBS (Gibco, Life Technologies Limited, USA) and 1% of Penicillin (10,000 units/mL)/Streptomycin (10,000 μg/mL)-solution (Gibco, Life Technologies Limited, USA) and one third of growth medium for Keratinocytes (Gibco, Life Technologies Limited, USA) supplemented with Bovine Pituitary Extract (25 mg) and EGF Human Recombinant (2.5 μg) (Gibco, Life Technologies Limited, USA). This medium was established through previous successes in obtaining cell lines from primary tissue [26]. Culture medium was changed twice a week. Cells were passaged in a dilution of 1:6 once they grew confluent.

### 2.5 Microscopy

Growth pattern and cell morphology was determined by phase contrast images using a Zeiss phase-contrast microscope Axio, Vert.A1 fitted with Axiocam 105 Color camera (Zeiss, Jena, Germany).

### 2.6 Protein isolation and quantification

Cells were lysed in a buffer containing 20 mM Tris-HCl, 150 mM NaCl, 5 mM EDTA, 1% Triton X-100, 0.5% sodium deoxycholate, 1 μM dithiothreitol (DTT), and proteinase and phosphatase inhibitors. Protein concentrations were measured using Coomassie-Reagents (Thermo Scientific, Rockford, USA) as per manufacturer’s instructions.

### 2.7 Western Blot

Western blots were performed as described in [31]. Briefly, 15 μg total protein per sample were loaded on a 10% SDS-polyacrylamide gel and subjected to electrophoresis. The separated proteins were transferred to a PVDF membrane (1 mA per cm^2^ at room temperature via semidry blot for 1 hour). Afterwards the blots were blocked in TBS containing 0.1% Tween 20 and 5% BSA at room temperature for 1 hour. Membranes were then probed with respective antibodies overnight at 4°C in a dilution as described in Table 2.

**Table 2:** Table describing the anti-bodies used in the study.

| Antibody | Dilution | Product no. | Company |
| --- | --- | --- | --- |
| Beta-Actin | 1:1000 | 4970S | Cell Signaling Technology |
| Keratin 19 | 1:1000 | 4558S | Cell Signaling Technology |
| P53 | 1:1000 | 48818S | Cell Signaling Technology |
| SMAD4 | 1:1000 | 38454S | Cell Signaling Technology |
| Anti-Rabbit - HRP conjugate | 1:1000 | 7074S | Cell Signaling Technology |
| Anti-Mouse - HRP conjugate | 1:1000 | 58802S | Cell Signaling Technology |

Equal protein loading was confirmed by stripping and re-probing the membrane with a monoclonal anti-human-β-actin antibody. For determining the level of mechanical stress on the cells, the Apoptosis Western Blot Cocktail (Abcam, USA) including pro/p17-caspase-3, cleaved PARP1 and muscle actin was used as per the manufacturer’s guideline. Detection of immunoreactive bands was performed using an ECL system (PerkinElmer, Inc., Netherlands).

### 2.8 Statistical Analysis

Figure layouts and graphs were prepared in with Graph Pad Prism 6.Ver.

## 3 Results

### TissueGrinder enables isolation of cells from various tumor tissues

Initially, 8 surgically resected tumor tissues from gastrointestinal (GI) tract were processed and an optimised protocol to isolate cells from resected tumor tissue for TissueGrinder was obtained Figure 3 and Table 3.

**Figure 3:**
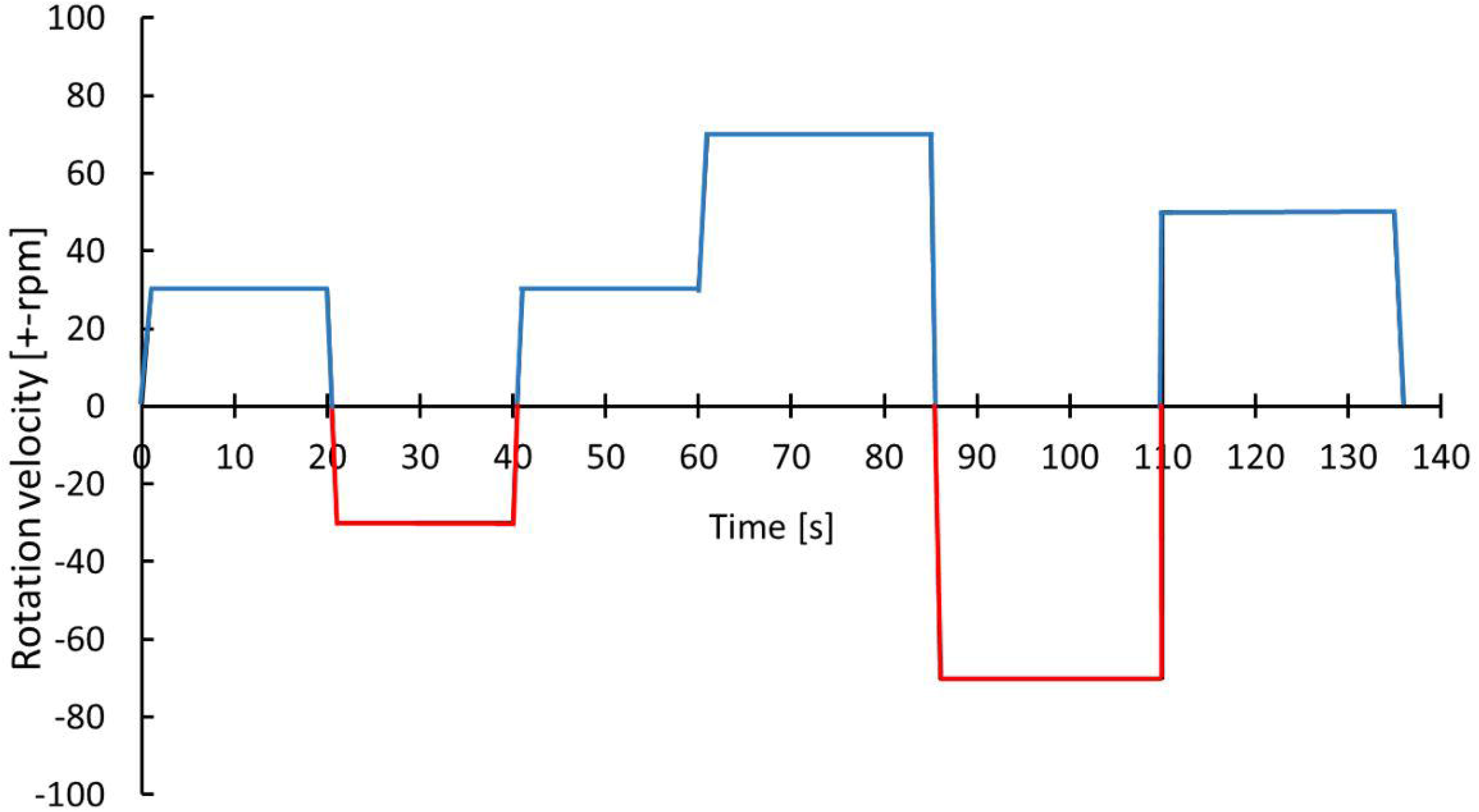
Optimized protocol for dissociating cells from GI track tumor tissue. With the blue color indicates the cutting while the red color indicates grinding process. The exact parameters and duration is indicated by alternations in rotational speed and direction over process duration.

**Table 3.** The parameters of the optimized TissueGrinder program for cell isolation.

| Steps | Program | Soft |  |
| --- | --- | --- | --- |
|  | Process | Speed<br>(rpm) | Duration<br>(sec) |
| 1 | Cutting | + 30 | 20 |
| 2 | Grinding | - 30 | 20 |
| 3 | Cutting | + 30 | 20 |
| 4 | Cutting | + 70 | 25 |
| 5 | Grinding | - 70 | 25 |
| 6 | Cutting | + 50 | 25 |

This protocol was then used for processing of 25 surgically resected tumor samples collected between October 2019 and June 2020. Table 4 mentions all the tumor samples used in this study. These samples were processed with TissueGrinder and explant method both to evaluate - ability to isolate living cells from various tumor tissues. These processed tumor samples were mainly from the pancreas, liver and colon-rectal regions. Some rare tumors entities like bile duct cancer (CCC-32), colon cancer (MCC-191), anal cancer, endometrial cancer metastasis, and gastric cancer (GCM-2) were also processed (Figure 4 A). Depending on the total amount of the tissue sample received from the surgically resected tumor, between 100 mg and 200 mg of the tissue was processed for cell isolation per grinding set per run. However, this was not possible for all the runs. 20 mg of tumor from pancreatic cancer (MaPaC-176) was the smallest amount of tissue processed while 305 mg from Liver cancer (HCC-184) was the largest amount of tumor tissue processed with TissueGrinder in a single run, which both yielded living cells. The range of the samples amount used for each cancer type is displayed in Figure 4 B. The most effective processing and cell yield was obtained when the processed tissue weighted between 100 mg and 200 mg.

**Table 4:**
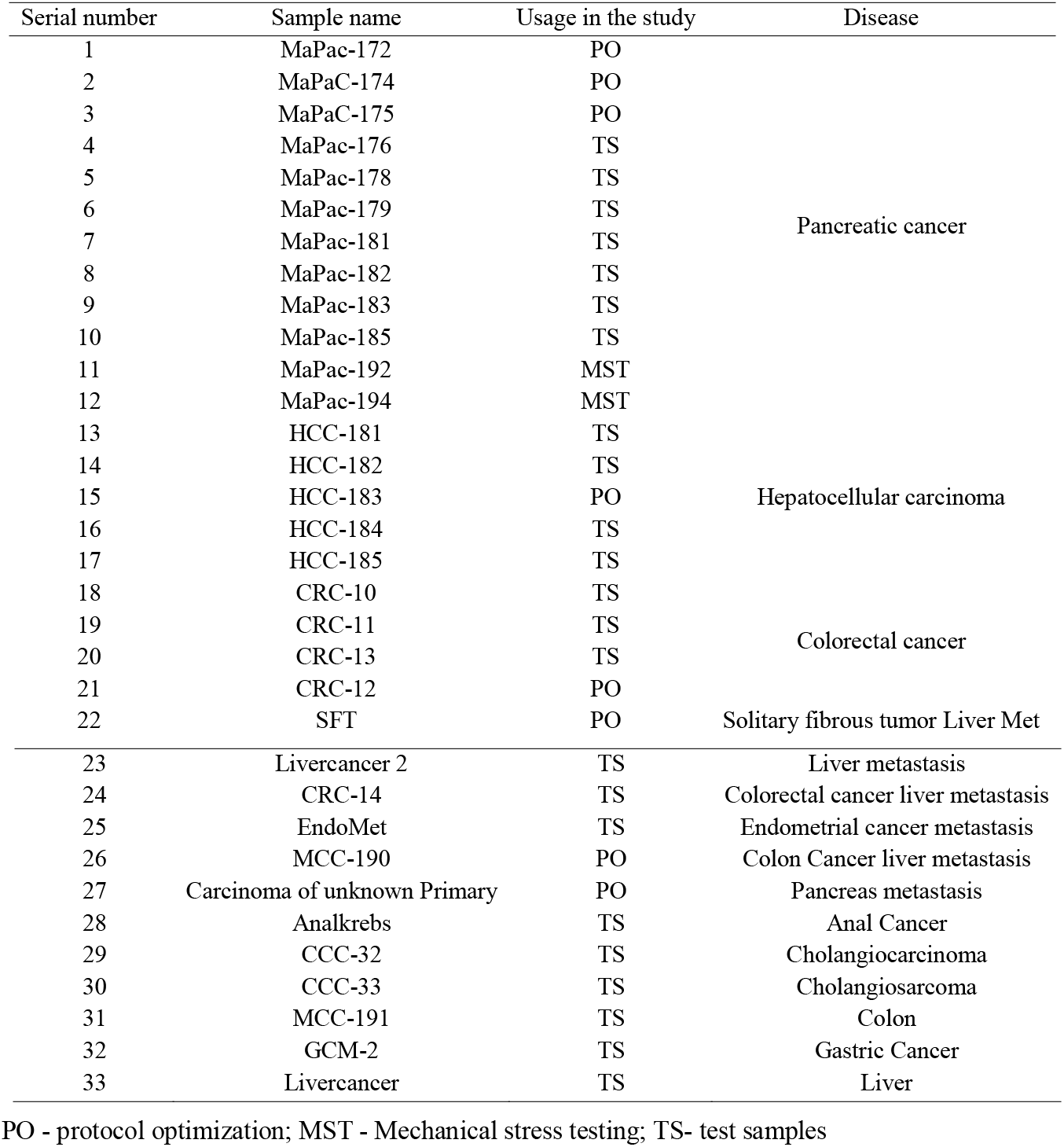
List of all the tumor samples processed in the study.

**Figure 4:**
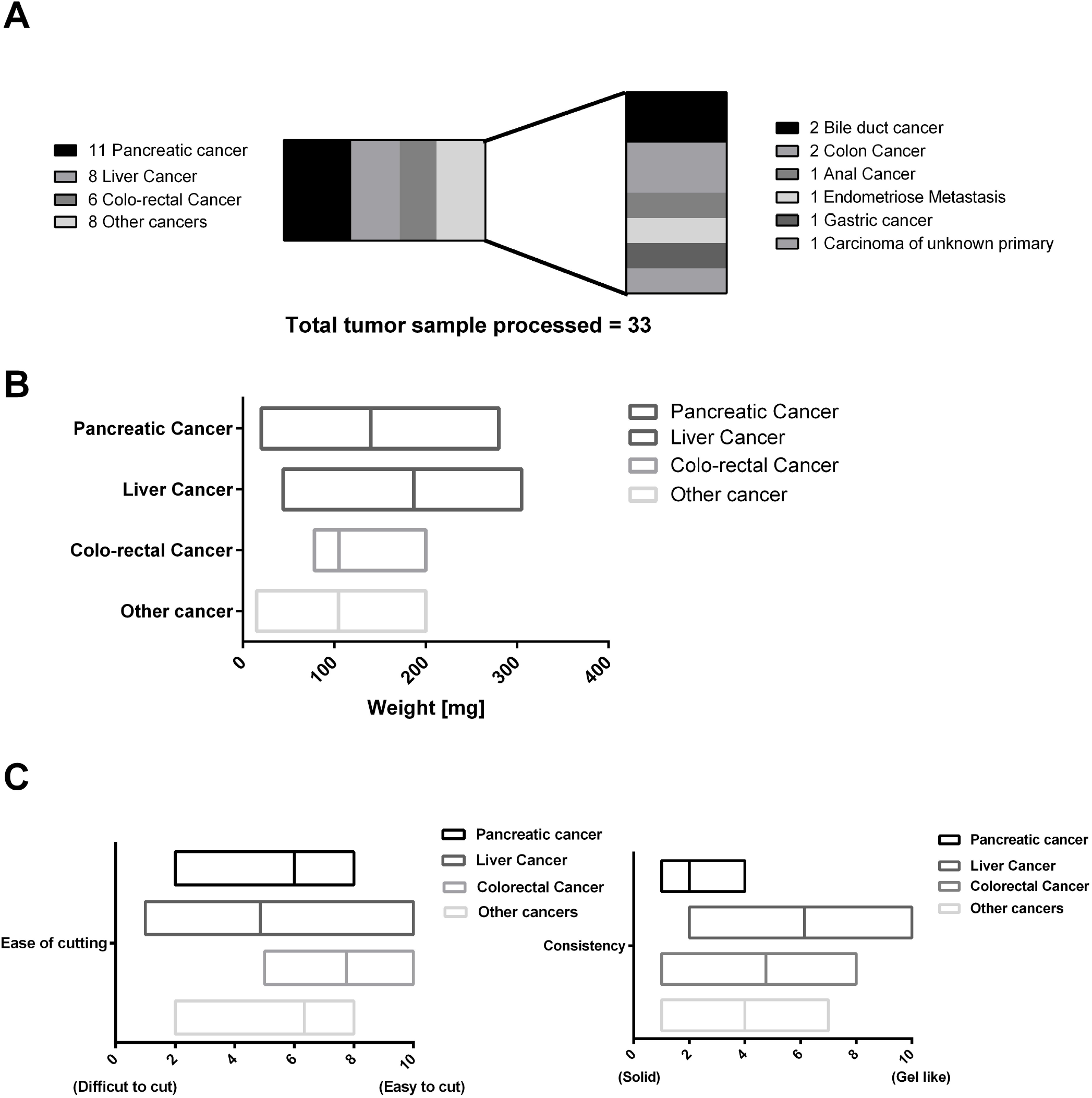
Overview of processed tumor samples using the TissueGrinder. A) 33 Tumor samples processed with the TissueGrinder and their categorization. B) The weight distribution of the tumor samples processed in each separate run for each of the four major tumor tissue categories with the median, minimum and maximum amount processed per run. C) The physical/material properties of the tissues processed based on ease of cutting and consistency. For ease of cutting ‘0’ means difficult to cut while ‘10’ represents easy to cut tissue. Consistency defined texture of the tissue processed where ‘10’ meant soft, gel like tissue while ‘0’ meant firm tissue.

Different types of tumor possess different physical and biological properties. The physical properties of the tissue processed in this study is defined based on a subjective scale where two properties – ease of cutting, and consistency of the processed tissue - are ranked on a scale of 0 to 10. Ease of cutting is described as – the ease with which tumor tissue was diced with the sterile scalpel during the sample preparation step, easy to cut samples were placed higher on the scale with 10 being most easy while difficult to cut samples were placed lower on the scale, with 0 being most difficult to cut tissue. Fibrotic and calcified tissues were difficult to cut compared to tumors which were neither fibrotic nor calcified. Consistency of the received sample was described on the basis of the firmness/texture of the sample observed macroscopically while cutting the tissue. Samples which displayed gel like or gooey properties were placed on the higher end (toward 10) and firm samples were placed on the lower end (towards 0). Some of the tumor samples like most liver cancer were mucigenous compared for example to most tumor samples obtained from pancreatic cancer which were more firm. Figure 4 displays this approach to categorize the variety of the processed tissue processed in this study.

### Tumor tissue processed with TissueGrinder yielded heterogeneous cells in shorter period of time compared to conventional explant method

One month post processing of the resected tissues samples, viable living cell were obtained in 75% of the samples processed with the TissueGrinder while only in 45% of samples processed with explant method living cells were observed, even though tissue from the same resected tumor sample and same amount of tissue was processed with both methods. Additionally, more cells have been observed in TissueGrinder processed samples compared to explant method. In many cases, more heterogenous cell populations from tumor were observed in TissueGrinder when compared to the explant method. An example is shown for visceral sarcoma, pancreatic cancer and anal cancer at 8, 20 and 47 days (Figure 5).

**Figure 5:**
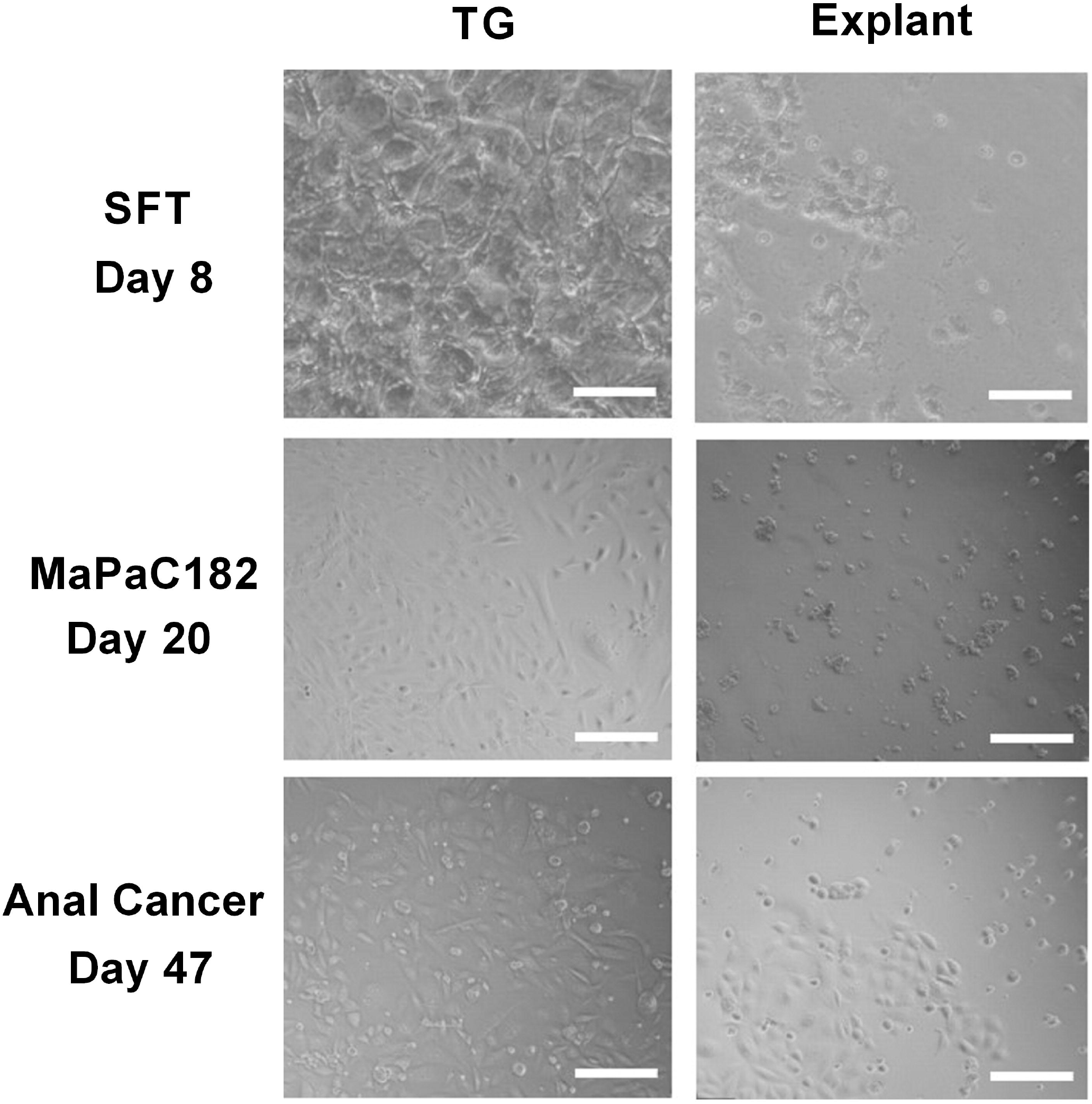
Comparison between three different tumor tissues processed with both TissueGrinder and explant method. Bright-field images of the tumor sample from sarcoma, pancreas cancer and anal cancer processed with TissueGrinder and explant method on 8th, 20th and 47th day after processing are shown. The images were taken with 4x Zeiss objective (LDA-Plan421231-991) The white scale bar represents 200 μm.

### Expression of specific markers for pancreatic cancer in three pancreatic cancer samples

To determine if the cells isolated from the tumor tissue were cancerous, total protein isolated from three subsequent pancreatic cancer samples processed by both explant and TissueGrinder and cultured for a period of six weeks was probed for the expression of pancreatic cancer markers via western blot. To this end expression of SMAD4, p53 and CK19 was evaluated. SMAD4 was observed in all cells irrespective of how the tumor sample was processed. Stronger expression of SMAD4 was observed in the MaPaC-181 and MaPaC-182 cells isolated via explant, however the MaPac-183 cells isolated with the TissueGrinder showed a reversed trend. The presence of p53 was observed in all of the tested samples irrespective of the method of isolation. CK19 seems to be just expressed in the explant sample of MaPaC-181 and the TissueGrinder sample of MaPaC-182. No expression of CK19 was observed in the MaPaC-183 cells.

### Mechanical stress endured by cells during isolation with TissueGrinder does not lead to Apoptosis

TissueGrinder employs alternating processes of cutting and grinding to isolate cells from diced tumor tissue. Although cells in tissue are generally exposed to multiple mechanical stresses during development, tissue homoeostasis and diseases yet excessive mechanical force can lead to cell death. Therefore, we further investigated if cells isolated via TissueGrinder exhibit Apoptosis via immunoblot. Total protein from cells isolated with TissueGrinder collected immediately, 24 h and 48 h after the processing of two pancreatic cancer tissue samples - MaPac192 and MaPaC194 from two different donors hardly showed cleaved caspase□3 (Figure 7). However, a slight expression of apoptosis-specific 89 kDa PARP fragment was observed in MaPaC 192 sample at all the time points, while its expression in MaPaC 194 was comparably lower and increased after 48 hours.

**Figure 6:**
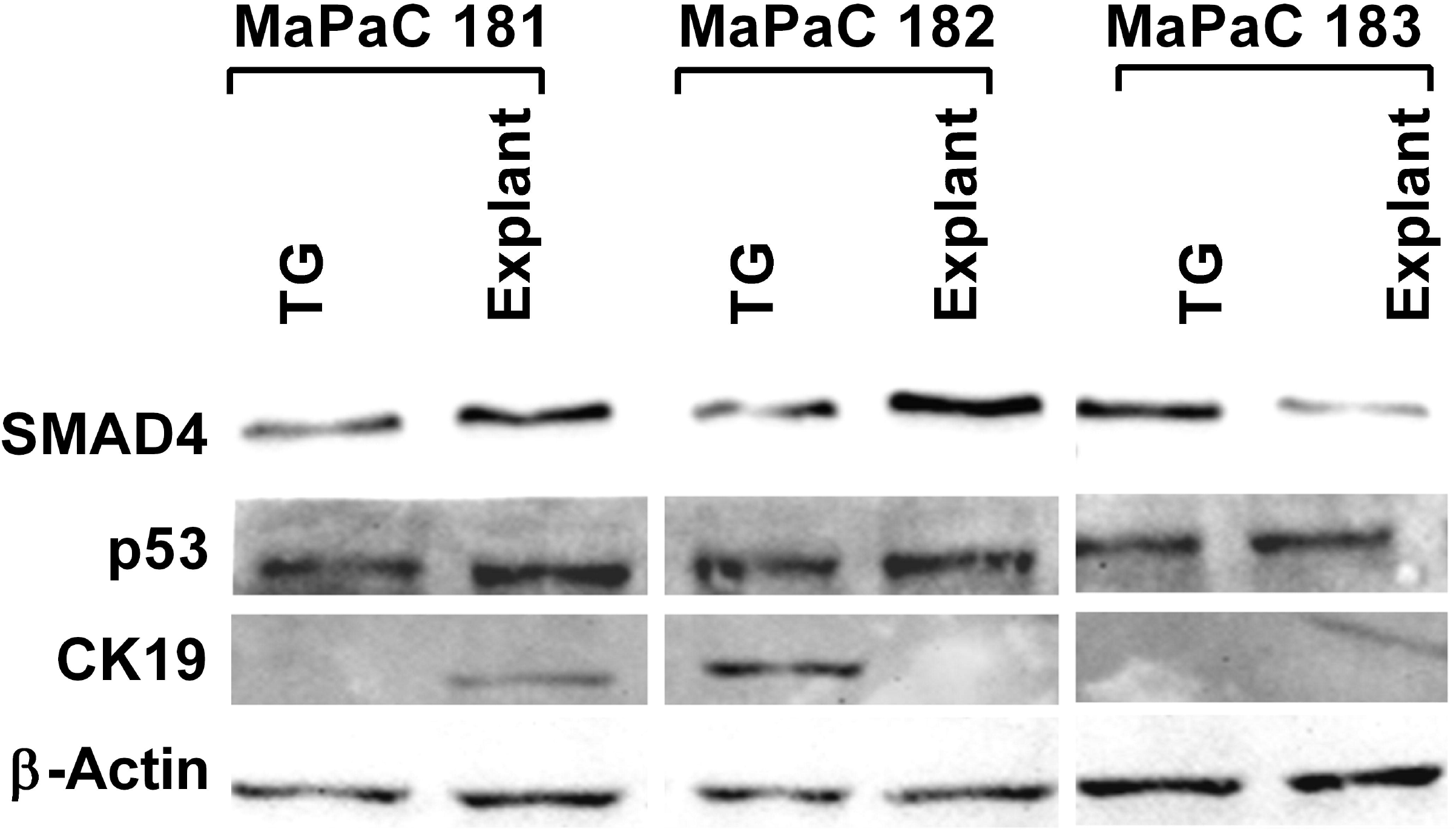
Expression of pancreatic cancer markers in three processed pancreatic cancer samples. Western blots were performed on the total protein isolated from cell obtained after processing three pancreatic cancer sample with TissueGrinder, explant method six weeks after the processing. Expression of the proteins SMAD 4, p53, and CK19 was evaluated. 20 g of total protein was loaded in each lane and β-Actin served as a loading control.

**Figure 7:**
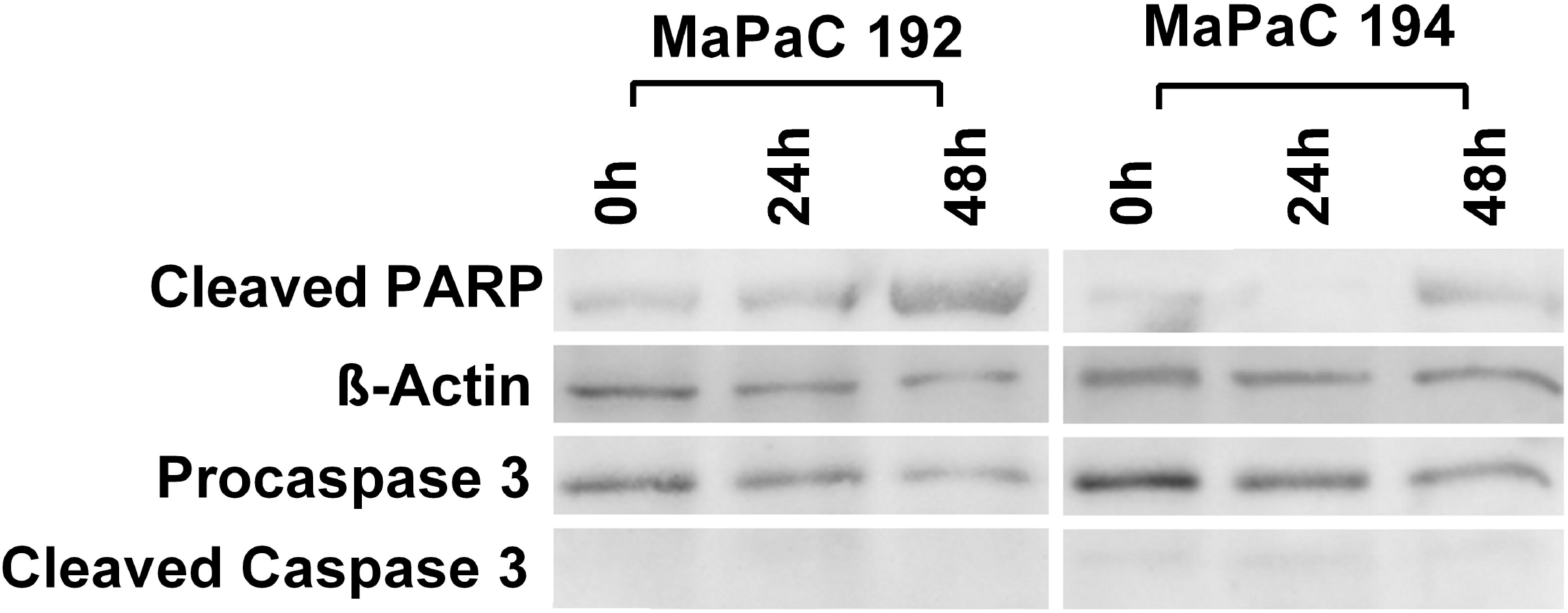
Mechanical stress endured by cells during processing by TissueGrinder does not lead to apoptosis. Western blots were performed on the total protein isolated from cell obtained after processing of pancreatic cancer tissue samples - MaPac192 and MaPaC194 from two different donors using TissueGrinder. The cells were collected immediately, 24 h and 48 h after the processing of the tissue. Expression of PARP (cleaved-PARP), caspase3 and cleaved caspase3 was investigated. 20 μg of total protein was loaded in each lane and equal protein loading was confirmed with ß-Actin.

## 4 Discussion

In this study firstly, we establish an application of the mechanical tissue dissociation device TissueGrinder to isolate primary tumor cells from a variety of tumor tissues. Secondly, we show that TissueGrinder can be used to isolate heterogeneous cell populations without introducing a bias for one cell type. Thirdly, we establish that mechanical force endured by cells during mechanical isolation in the TissueGrinder device are not leading to apoptosis in the isolated cells. Fourthly, we report that primary tumor cells were obtained a week earlier when TissueGrinder was used in the comparison to the explant method.

The first step in isolating individual cells from primary samples is to produce a suspension of viable single cells. This is not trivial when working with complex solid tissues −the extracellular matrix, which holds the different cell types together, varies drastically in its composition from one tissue type to another [32]. Due to its highly adhesive characteristics, it ensures that the cells are held together and can thus function as a closed network. Furthermore, diseased tissues can have different dissociation kinetics when compared with their normal counterparts, as well as varied dissociation between samples of the same disease [23]. For instance fibrosis and calcification are two common processes found in several kinds of tumors which impede mechanical dissociation of tissue. Thus, a delicate balance between dissociating the tissue and preserving the cells is challenging but required [25]. With the possibility to program the grinding and cutting processes in TissueGrinder, individual processing profiles for specific tissues were generated and cells were obtained from various cancer tissue samples.

Tumors belong to a complex entity and consist of various cells which interact with each other. For instance, tumor fibroblasts and tumor cells are implicated in chemoresistance [33, 34]. Therefore, it is important that cell heterogeneity is maintained in primary cells isolated from tumor tissue to determine for example effectiveness of chemotherapeutic drugs. Cells obtained after processing with the TissueGrinder were often more heterogeneous compared to these obtained via explant method. In explant method, cells mostly fibroblast cells grow out of the tissue into the culture dish. Even though the tissue is diced into a small piece of 1 mm^3^ yet there is a gradient of nutrition, with less amount reaching the cells in the core of the cut tissue piece. As a result, only certain cells are able to grow out of the tissue. Contrary to that, diced tissue is further cut and grinded into smaller fragments by the TissueGrinder and cells are released into harvesting medium. Therefore, cells isolated by TissueGrinder retain the tumor heterogeneity better when compared to the cells isolated via explant process.

Each method of sample preparation, i.e. cell isolation from tissue, could have a different degree of adverse effects on cell integrity post processing, which should be minimized as much as possible. Typical stress factors are - suboptimal temperature, suboptimal buffer conditions, stress caused by long enzymatic dissociation or high mechanical stress during grinding or damage caused by centrifugation, resuspension or vortexing of cells. The effects on the cells often depend on the time of exposure to these factors. With TissueGrinder the possibility to program the instrument helps to minimize these factors as much as possible to achieve a gentle dissociation protocol. This is clearly evident as tissues processed via TissueGrinder are subjected to mild mechanical stress slight expression of cleaved PARP-1 yet it does not lead to apoptosis the absence of cleaved caspase-3.

Given the increasing number of samples in medical, pharmaceutical and biological analysis and the fact that sample preparation is still one of the most time-consuming processes, the need for more efficient approaches is increasing. Furthermore, downstream molecular biological analyses require a high degree of sterility and molecular integrity of the target cells. Cell integrity is often required throughout the cell isolation process. In particular, if the genome or proteome is the target of analysis, cell integrity should be maintained prior to lysis to avoid early degradation of DNA/RNA. Cell viability is required when single cells are isolated for the production of cell cultures or to study stem cell differentiation. Cells react to stress factors such as mechanical forces, radiation, chemical changes in their environment, etc., which can lead to differentiation, reduced viability or even apoptosis. Technologies should provide sufficient “gentle” extraction and handling when working on living cells. TissueGrinder addresses this issue with shortening time to get cells but also through reduced handling errors and increased reproducibility. As no enzymes are employed in the dissociation of the cells with TissueGrinder, cells are as close as currently possible to their native environment. Thus, the cells isolated via TissueGrinder can be utilized for various analytical applications like FACS sorting, NGS, mass spectrometery and drug screening. Although the tissue dissociation workflows prior to the actual analysis are still a time-consuming and they are not completely reproducible as well, since they are also performed manually [15]. Compared to the traditional method of cell isolation via explant method, it takes around 15 minutes – 30 minutes longer to process tissue using the TissueGrinder yet cells have been observed sooner in the TissueGrinder cultures (after 4-7 days) than in the explant cultures (after 10-12 days). Therefore, samples processed by TissueGrinder were nearly one week earlier available for cell analysis than the ones processed with explant method.

The possibility of extracting individual cells from tissues is currently a bottleneck for cell-based diagnostic technologies and remains a decisive factor in the fields of personalized medicine. Manual processing of tumor sample makes reproducibility of results difficult. Preliminary dissociation methods with a continuous workflow in a closed system provides advantage of avoiding cross-contamination and manual errors. Although the TissueGrinder was used in this study in semi-automatic manner, the modular design of TissueGrinder will easily facilitate automation. Furthermore, in the development of mechanical tissue dissociation processes, it is not possible to rely on theoretical material values due to the wide range of samples and the corresponding complexity of the tissue properties. For an automated tissue dissociation, it is therefore necessary to provide specific protocols for different tissues. The possibility to program the TissueGrinder not only yields a time benefit but also facilitates generation of tissue specific isolation protocols of cells from specific tumor types.

The massive impact of personalized medicine approach is currently limited by a complex process control, missing interface to common laboratory devices and time-consuming individual sample preparations leading to a very limited sample throughput and rather demanding requirements on operator training. The possibility of extracting individual cells from tissues is currently a bottleneck for cell-based diagnostic technologies and remains a decisive factor in the fields of personalized medicine. TissueGrinder addresses these issues with shortening not only the time of processing but also reduced handling errors and increases reproducibility. As no enzymes are employed in the dissociation of the cells with TissueGrinder, cells are as close to their native environment. Thus, the cells isolated via TissueGrinder can be utilized for various analytical applications like FACS sorting, NGS, Mass-spectrometery and drugs-screening.

### Limitations

Although in this paper we developed a general protocol for isolation of the cells from primary solid tumors, the protocol can be further optimized for specific tumor tissues. Therefore, this protocol should be taken as basis and can be further optimization for criteria such as viability, and yield of cells of interest. In this study we did not use yield i.e. number of cells isolated per mg of tissue as parameter for optimization as tumor is a complex tissue which is infiltrated with various immune cells and many times in the core of the solid tumor there are dead cells due to hypoxia and nutrition gradient. Therefore, number of cells isolated per mg of tissue, i.e. yield might not be ideal approach for isolation of cells from tumor tissue. Furthermore, Anoikis induction of apoptosis in cells upon loss of attachment to the extracellular matrix (ECM) and neighboring cells could also result in low yield. Therefore, use of Anoikis inhibitors for purpose of increasing cell viability could also be considered.

## 5 Conclusion

TissueGrinder is a novel technology for rapid generation of patient-derived single cell. It can be adapted to various tissue types isolate various cells of interest and provides reproducible results. This technology can be integrated in any personalized medicine workflow.

## 6 Conflict of Interest

SS and JL are co-founder of Fast Forward Discoveries GmbH distributing TissueGrinder-Technology. All other authors do not report a conflict of interest.

## 7 Ethics statement

The tumor tissue used for the isolation of cells received from donors operated at Department of Surgery, Medical Faculty Mannheim, Heidelberg University, Mannheim, Germany. This was approved by the local ethics committee (Medizinische Ethikkommission II der Medizinischen Fakulta□t Mannheim), and all of the donors gave their written, informed consent (Ethics Application No. 012-293N-MA).

## 8 Author Contributions

RRM, SS, and PP conceived and planned the experiments. RRM, JML and PP carried out the experiments. FR and CR contributed to sample preparation. JML, RRM, SS, JL, FR and PP contributed to the interpretation of the results. SS, RRM and PP wrote the manuscript. All authors provided critical feedback and helped shape the research, analysis and manuscript.

## 9 Funding

This research work was supported by Wilhelm Müller-Stiftung, granted to FR and PP.

